# Functional Analysis of Distinct Populations of Subthalamic Nucleus Neurons on Parkinson’s Disease and OCD-like Behaviors in Mice

**DOI:** 10.1101/2020.06.10.137679

**Authors:** Luca Parolari, Marc Schneeberger, Nathaniel Heintz, Jeffrey M. Friedman

**Affiliations:** Laboratory of Molecular Genetics, Howard Hughes Medical Institute, The Rockefeller University, New York, NY 10065, USA; Laboratory of Molecular Biology, Howard Hughes Medical Institute, The Rockefeller University, New York, NY 10065, USA

## Abstract

The Subthalamic Nucleus (STN) is a component of the basal ganglia and plays a key role to control movement and limbic-associative functions. STN modulation with Deep Brain Stimulation (DBS) improves the symptoms of Parkinson’s Disease (PD) and Obsessive-Compulsive Disorder (OCD) patients. However, DBS does not allow for cell-type specific modulation of the STN. While extensive work has focused on understanding STN functionality, the understanding of its cellular components is limited.

Here, we first performed an anatomical characterization of molecular markers for specific STN neurons. These studies revealed that most STN neurons express *Pitx2*, and that different overlapping subsets express *Gabrr3*, *Ndnf* or *Nos1*. Next, we used neuronal modulatory tools to demonstrate their roles in regulating locomotor and limbic functions in mice. Specifically, we showed that optogenetic photoactivation of STN neurons in Pitx2-Cre mice or of the Gabrr3-expressing subpopulation induces locomotor changes, and improves locomotion in a PD mouse model. Additionally, photoactivation of Pitx2 and Gabrr3 cells induced repetitive grooming, a phenotype associated with OCD. Repeated stimulation prompted a persistent increase in grooming that could be reversed by fluoxetine treatment, a first-line drug therapy for OCD. Conversely, repeated inhibition of STN^Gabrr3^ neurons suppressed grooming in Sapap3-KO mice, a model for OCD. Finally, circuit and functional mapping of STN^Gabrr3^ neurons showed that these effects are mediated via projections to the globus pallidus/entopeduncular nucleus and substantia nigra reticulata. Altogether, these data identify Gabrr3 neurons as a key population in mediating the beneficial effects of STN modulation thus providing a new molecular handle for PD and OCD drug discovery.

## INTRODUCTION

Parkinson’s Disease (PD) and Obsessive-Compulsive Disorder (OCD) are highly debilitating neuropsychiatric disorders (1,2) associated with alterations in dopaminergic signaling (3,4). Affected patients benefit from Deep Brain Stimulation (DBS) when pharmacologic and behavioral therapies prove inadequate (5). DBS consists of a permanent neurosurgical device that delivers electrical current in specific brain regions, including most frequently the Subthalamic Nucleus (STN) (6–9).

The STN is a crucial node within the indirect pathway of the basal ganglia and also receives significant hyper-direct frontal cortex inputs. STN neurons are thought to inhibit movement initiation via excitatory projections to the globus pallidus interna and substantia nigra reticulata. In a physiologic state, the activity of the STN is suppressed by dopaminergic signaling. In diseases when dopaminergic signaling is abrogated such as PD, this nucleus is hyperactive and thus blocks movement (10,11).

Importantly, it has been hypothesized that DBS’s high-frequency electrical current improves PD symptoms by decreasing the activity of STN neurons (12). Consistent with this hypothesis, anatomical lesions of the STN show similar effects to DBS in patients (6,13) and non-human primates (14). Additionally, growing evidence has implicated the STN in emotion regulation and associative functions (15). Several clinical trials have confirmed the efficacy of DBS-STN as a treatment for OCD (8,9), but have also reported that DBS-STN causes acute (16) and chronic episodes of depression, and impairments in cognitive functions and impulse control (17–19). Similarly, pharmacologic hyperactivation of the STN in monkeys can induce an array of behaviors, including compulsive, stereotyped behaviors characteristic of OCD such as repetitive grooming, smacking, and licking (20). In rodents, high-frequency stimulation of the STN can suppress excessive self-grooming in autism-like mouse models (21) and reduce compulsive lever pressing in rats (22), while STN lesions impair impulse control (23,24). The STN has also been involved in the reward circuit, both in evaluating the valence of stimuli (reward/aversion) (25,26) and of action outcome (27). However, while it is understood that the STN plays a crucial role in several pathologic conditions, the neural populations responsible for these effects and their function in the basal ganglia circuit remain largely unknown.

Here we provide an anatomic characterization of different STN neuronal subtypes, identifying three clusters defined by the markers *Nos1*, *Ndnf* and *Gabrr3*, a subunit of GABA-A receptor expressed at low levels in brain. Functional studies revealed that increasing the activity of the STN and its Gabrr3 subpopulation using optogenetics can induce both PD- and OCD-like phenotypes. Modulation of these neurons is also effective in reversing the pathologic phenotypes in animal models of these conditions. Finally, circuit and functional dissection of these neuronal populations show that these behavioral effects are mediated via projections to the globus pallidus/entopeduncular nucleus (GP/EP) and substantia nigra reticulata (SNr).

## METHODS

### Multiplex Fluorescence In Situ Hybridization (FISH)

Mice were anesthetized and perfused transcardially with RNase-free PBS, followed by 4% PFA. Brains were harvested and post-fixed overnight in 4% PFA, incubated for 24h sequentially in 10%, 20% and 30% glucose solution, sliced with temperature-controlled cryostat (17um thickness), and mounted on Superfrost Plus Adhesion slides (Fisher). Multiplex FISH was then performed using the RNAscope system (ACDBio) as per the manufacturer’s protocol. The slides were mounted using Prolong Gold Antifade Mountant (Thermo Fisher). The target probe sets used included vGlut2 (Slc17a6), vGat (Slc32a1), NeuN, Pitx2, Nos1, Ndnf and Gabrr3. Images were captured using confocal microscopy (Zeiss or Leica), and subsequently manually counted for the number of cells single-, double- or triple-labelled fluorescent cells using Fiji ImageJ.

### Animals

All experiments were conducted in accordance with the guidelines of the National Institutes and the protocols approved by the The Rockefeller University Institutional Animal Care and Use Committees. Mice were housed in a 12 hr light-dark cycle (lights on at 7:00) with ad libitum access to food and water. Male mice were used for all the behavioral and histological studies; both male and female mice were used for tracing studies. Mice were at least 8 weeks old at the time of surgery. All mouse lines were in a wild-type (C57BL/6J; Jackson Laboratory; stock 000664) background. The following mouse lines were used: Pitx2-Cre (Dr Tim Cox, The University of Washington (28)), Gabrr3-Cre (KC112 line; GENSAT; Dr Nat Heintz, The Rockefeller University), Sapap3-KO (Jackson Laboratory; stock 008733). For all experiments, hemyzygous Cre-positive animals were compared with wild-type littermates as controls. For Sapap3-KO experiments, knockout animals were compared with heterozygous littermates. A het × KO breeding scheme was used to yield a sufficient number of knockouts with heterozygous littermates as controls.

### Viral Vectors

All viruses used in these experiments were purchased from Addgene and UNC Vector Core. The following viruses were used for optogenetic studies: AAV5-EF1a-DIO-hChR2(H134R)-EYFP (activation; Addgene 20298), AAV5-EF1a-DIO-eArch3.0-eYFP (inhibition; UNC Vector Core). For tracing studies: AAV5-hSyn-DIO-hM3D(Gq)-mCherry (anterograde; Addgene 44361), rgAAV-hSyn-DIO-EGFP (retrograte green; Addgene 50457), rgAAV-CAG-FLEX-tdTomato (retrograde red; Addgene 28306). For ablation study: AAV8-EF1a-lox-Cherry-lox-(dtA)r-lox2.ape (lot # AV6241B).

### Stereotaxic surgeries

Mice were anesthetized using isoflurane. Coordinates were identified using the Paxinos mouse brain atlas and adjusted over several rounds of practice usign flurescent beads. For viral injections (1.0uL Syringe, Hamilton 65458-01), the coordinates used were: STN (ML: ±1.68mm, AP: −2.00mm, DV: −4.68mm), GP/EP (ML: ±1.85mm, AP: −1.15mm, DV: −4.10mm), SNr (ML: ±1.50mm, AP: −3.80mm, DV: −4.20mm). Viral injection volumes were STN (0.04uL), GP/EP (0.10uL), SNr (0.30uL). Fiber optic ferrule (Thor Labs) implants were positioned at the same ML and AP coordinates, and 0.40mm more dorsal than the injection site (ie, STN (DV: −4.28mm), GP/EP (DV: −3.70mm), SNr (DV: −3.80mm) to avoid structural damage to the targeted area. All listed DV coordinates are relative to pia. Optic fibers were seecured in place with dental cement. Viral injections and optic fiber implants were performed unilaterally or bilaterally as specificed in the Results section. Skin was closed using 5-0 silk sutures.

Considering the small size of the targets in mice, in particular the STN, each animal’s skull was carefully adjusted to lay flat, allowing for a skew of 0.01mm or less both along the AP axis (ie, delta DV between bregma and lambda at skull surface) and ML axis (delta DV between ML: +1.68mm and ML: −1.68mm, at AP: −2.00mm at skull surface). Of note, AP coordinates for the STN were adjusted (AP −2.00mm ± 0.15mm) depending on the size of each animal skull, measured as distance between bregma and lambda.

To produce hemiparkinsonian mice, animals were injected with 20mg/kg i.p. desipramine hydrochloride (Sigma, D3900) 30-60 minutes prior to surgery to prevent 6OHDA-mediated noradrenergic lesion; 30-60 minutes later, animals were injected in the medial forebrain bundle (ML: ±1.20mm, AP: −1.10mm, DV: −5.20mm) with 0.2 mg/kg 6OHDA hydrbromide (Sigma, H116) dissolved in 0.1% ascorbic acid. To assist during post-op recovery, animals were i.p. injected daily with dextrose-saline solution and provided condensed milk *at libitum* for 5 days, as well as checked for their health condition daily until day 10 post-op.

### Behavioral Assays

#### Optogenetic Induction of Rotation Behavior in Healthy and Hemi-Parkinsonian Mice

Pitx2-Cre, Gabrr3-Cre and control littermate mice were injected with channelrhodopsin or archaerhodopsin in the STN, followed by 3 weeks to allow for opsin expression and animal recovery; in the case of optogenetic stimulation of STN projections, at least 6 weeks were left to allow robust expression of Channelrhodopsin in the terminals.

On the day of the experiment, animals were placed in a clean and empty cage during the light phase, and connected to the laser via rotary joint (ThorLabs). Mice received 3-minutes laser stimulation with continuous green (532nm) or intermittent blue light (473nm, OEM Lasers/OptoEngine) at frequencies of 10, 40 or 120 Hz, 10ms pulse width, 5-10mW. Stimulation paradigms were programmed into an arbitrary waveform generator (Agilent). In all experiements, mice were given *ad libitum* access to food prior to and after the assay. Video was recorded with Logitech Webcam for offline analysis for a total of 9 minutes (3 minutes before, during and after stimulation). Rotations were recorded using a hand tally counter (Fisher Scientific) as full-body turns of at least 180° in either direction. The rotation ratio was calculated as counter-clockwise divided by clockwise rotations if the ratio was ≥ 1; if this was < 1, then the ratio was reported as the opposite of its reciprocal. When the number of rotation performed in one direction was 0, the count was rounded to 1.

Hemiparkinsonian mice were left to recover from surgery for at least 6 weeks before the experiements; 30 minutes before the behavioral measurements, mice were i.p. injected with 2.6mg/kg of amphetamine hemisulfate (Sigma, A5880) to induce rotations ipsilateral to the 6-OHDA lesion. The subsequent behavioral paradigm was then carried out as detailed above.

#### Optogenetic Induction of Acute and Chronic Grooming and Reversal with Fluoxetine

Pitx2-Cre, Gabrr3-Cre and control littermate mice were injected with channelrhodopsin in the STN, followed by 3-6 weeks windows to allow for recorvery and opsin expression in the cell soma and terminals, respectively. For habituation to the assay, mice were placed in an empty shoebox cage, connected to the laser cable via their optic fiber implants, and left to get accustomed to the setting for 1 minute/day for 7 consecutive days. On the 8th day, during the light phase, each animal was placed in a clean and empty cage and connected to the laser cable bilaterally. After 3 minutes of habituation to the novel environment, mice were video recorded while freely moving in the cage for 5 minutes, followed by 5 minutes of optogenetic stimulation (40Hz, 10ms pulse, 5-10mW), and 5 minutes after the laser was turned off. The acute grooming was manually scored with JWatcher to quantify the time spent grooming and the number of grooming bouts during each interval.

To induce the persisten grooming phenotype, optimal 40Hz optogenetic stimulation was repeated for 5 minutes/day for 7 consecutive days, at a consistent our of the day. On the 8th day, mice were placed in a clean and empty cage and left to habituate for 3 minutes, plugged to the cable, recorded for 5 and the time spend grooming was quantified.

Mice then underwent an additional 14 consecutive days of daily 5-minutes optogenetic stimulations while treated with fluoxetine hydrochloride (Sigma, 1279804) 18mg/kg/day in drinking water, which was changed every 3 days to prevent inactivation of the drug. After 2 weeks of treatment, on day 21st, mice were videorecorded one last time for 5 minutes while connected to the cable. Mice were given *ad libitum* access to food and water prior, during and after the 21-day assay. Total locomotion was measured on the videos recorded using Noldus Ethovision XT 9.0 Software.

#### Optogenetic Reversal of Increased Grooming in Sapap3 KO mice

Gabrr3-Cre and control littermate mice, each divided in two groups of heterozygous and homozygous Sapap3 knock-outs, were bilaterally injected with archaerhodopsin and implanted with optic fibers, followed by a 3-week recovery window. After 7 days of habituation to handling and to the cable, mice were optogenetically inhibited with continuous green light for 10 minutes/day for 7 days. The total time spent grooming was measured with Home Cage Environment computerized scoring system before the 1st day, and 24h after the 7th day of daily optogenetic inhibition.

#### Computerized Scoring of Grooming with Home Cage Environment (Clever Sys Inc.)

Mice were transported in their housing cages to the procedure room 1h before testing for acclimation. 15 minutes before the experiment, animals were single-caged in clean new shoe-size cages, and the Home Cage Environment (Clever Sys Inc.) was calibrated for cage position and background. Recording was conducted for 2.5 hours, during which time the animals were left free to explore the cage, during light cycle, with ad libitum access to food and water. In the case of repeated measurements over different days (ie, before/after optogenetics or fluoxetine treatment) the same animals were recorded at consistent times during their light cycle.

### Immunohistochemistry

Mice were transcardially perfused with PBS, followed by 10% formalin. Brains were then post-fixed for 24-36h in formalin 10%, and sectioned by vibratome (50um thickness). Primary antibodies used for immunohistochemistry were chicken anti-GFP (Abcam, 13970), chicken anti-RFP (Abcam 62341), rabbit anti-RFP (Rockland, 600-401-379) and rat anti-TH (Immunogen, AB152); secondary antibodies were Alexa Fluor conjugated (Life Technologies). Slides were mounted using Prolong Gold Antifade Mountant (Thermo Fisher). All images were captured using confocal microscopy (Zeiss or Leica).

### Brain Clearing and iDISCO

Mice were perfused transcardially with PBS followed by 4% PFA. After a 24 hr post-fixing period, immunolabeling and whole-brain clearing and immunolabeling was performed (29). A LaVision Ultramiscrope was used for light sheet imaging of cleared brains. Antibody used was rat monoclonal anti-mCherry (16D7; ThermoFisher M11217). For acquisition, cleared samples were imaged in a sagittal orientation (left lateral side up) on a light-sheet microscope (LaVision Biotec) equipped with a sCMOS camera and LVMI-Fluor 4x objective lens equipped with a 6-mm working distance dipping cap. Version v144 and v210 of Inspector Microscope controller software was used. Samples were scanned in the 640 nm channel. Images were taken every 6 mm and reconstructed with Imaris 9.1 software for visualization. For autofluorescence, the 480 nm channel was used with a 1.3x objective lens.

### Quantification and Statistical Analysis

Statistical parameters reported are: sample size (n = number of animals or samples per group), mean, statistical test used, and statistical significance. All data are displayed as mean ± SEM. Significance was defined as p < 0.05. Significance annotations are: * p < 0.05, ** p < 0.01, *** p < 0.001, **** p < 0.0001. Mice were randomized into control or treatment groups. Control mice were age-matched littermate controls where possible. All statistics and data analysis were performed using GraphPad Prism 9.

## RESULTS

### Identification of Specific Markers of the STN and Its Subpopulations

The STN is comprised of mostly excitatory glutamatergic neurons defined by the marker *vGlut2* (30,31). However, the possibility that there are molecularly distinct subsets is still debated. To further identify specific molecular subpopulations of neurons in the STN, we first scrutinized in-situ hybridization (ISH) studies from the Allen Brain Atlas (32) and the Gene Expression Nervous System Atlas (33) databases. STN-specific genes from these experiments were scored and ranked based on STN expression intensity, density, local specificity, as well as overall central nervous system specificity (Supplementary Table 1). Intriguingly, four genes exhibited significantly high expression in the STN in comparison to neighboring areas and to the rest of the brain: Paired-Like Homeodomain 2 (*Pitx2*), Nitric Oxide Synthase 1 (*Nos1*), Neuron Derived Neurotrophic Factor (*Ndnf*), and Gamma-AminoButyric Acid Receptor Subunit Rho-3 (*Gabrr3*). To confirm their specificity to the STN we conducted brain-wide ISH studies in slices. These studies confirmed their expression in the STN, further suggesting that these markers might constitute specific molecular handles for STN neuromodulation (Figure 1A). In addition, we conducted quantitative co-localization measurements from the ISH studies to further characterize the identified markers for STN subpopulations. Consistent with a recent report, ISH for *Pitx2* showed extensive co-localization with fluorescent microscopy nuclear counterstain 4’,6-diamidino-2-phenylindole (DAPI) in the STN, suggesting that most STN cells express *Pitx2* (34). Furthermore, colocalization studies with the pan-neuronal marker neuronal-nuclei (*NeuN)* demonstrated that *Pitx2*-expressing cells are neurons (Figure 1B, left). Nearly all (>95%) *NeuN*-positive cells in the STN were co-labeled for *Pitx2*, thus *Pitx2* can be referred to as a general STN marker (hereafter referred to as STN^Pitx2^. Of note, a small number of cells were negative for both *NeuN* and *Pitx2* markers, suggesting the existence of a non-neuronal population of cells in the STN (Figure 1B, bottom left, white arrow). Next, we evaluated whether STN^Pitx2^ neurons are excitatory or inhibitory by coupling ISH studies for *Pitx2* and classical neurotransmitters. Importantly, ISH co-localization studies for *Pitx2* and either the glutamatergic marker *vGlut2* (*Slc17a7*) or the GABAergic marker *vGat* (*Slc32a1*) (Figure 1B, center and right) demonstrated that most STN^Pitx2^ neurons co-express *vGlut2*+, consistent with prior results (34), while none expressed *vGat.* However, we did note the presence of GABAergic neurons in the Zona Incerta (Z.I.), an adjacent structure.

**Figure 1.**
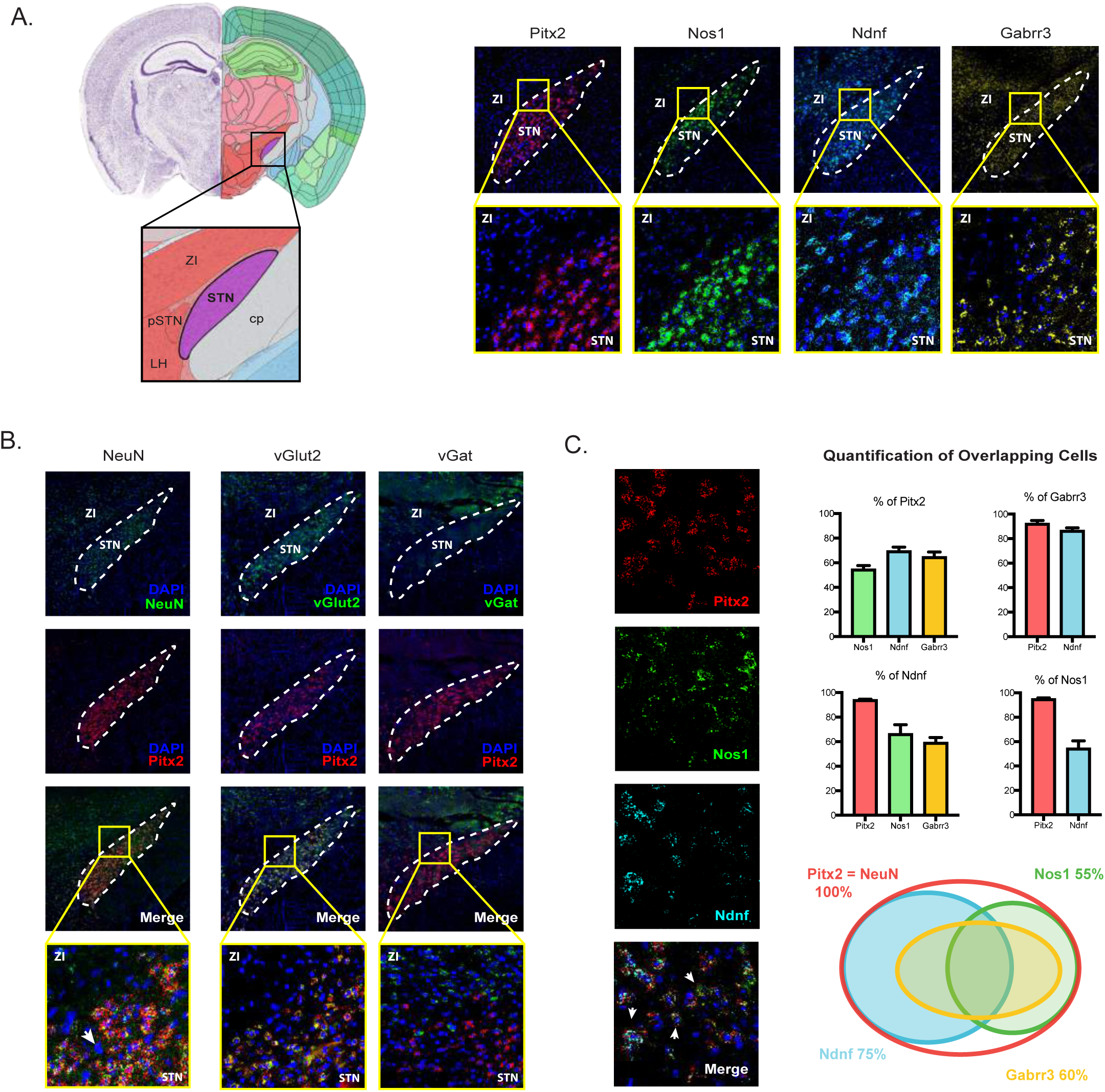
Identification of Subthalamic Nucleus (STN) Neuronal Subpopulations. (A) Left: schema showing the location of the STN in the mouse brain (coronal view at AP: −2.0mm from bregma). Right: fluorescence *in situ* hybridization (FISH) of the STN confirming the expression of genes *Pitx2*, *Nos1*, *Ndnf*, *Gabrr3*. Top row: whole view. Bottom row: 20x magnification of border between STN and ZI. STN: Subthalamic Nucleus; pSTN: Para-Subthalamic Nucleus; ZI: Zona Incerta; LH: Lateral Hypothalamus; cp: Cerebral Peduncle. (B) Characterization of *Pitx2* neurons by colocalization with neuronal marker *NeuN* (left), glutamatergic *vGlut2* (center) and GABAergic *vGat* (right). White arrow designates DAPI+, NeuN-Pitx2- non-neuronal cells. (C) Colocalization (and quantification) between *Pitx2*, *Nos1*, *Ndnf* and *Gabrr3* expressing-neurons. Left: example of triple FISH experiment showing colocalization between *Pitx2*, *Nos1* and *Ndnf* neurons; white arrows designate triple-labeled cells. Right, top: percentage of cells expressing gene on Y axis that coexpress gene on X axis. Data are presented as mean ± SEM. Right, bottom: Venn diagram summarizing quantitative overlapping of the 4 neuronal populations.

In contrast to *Pitx2*, *Nos1*, *Ndnf* and *Gabrr3* were expressed only in smaller clusters of STN cells. To further investigate how these markers are anatomically distributed, we next performed multiple fluorophore ISH experiments and quantified the extent of overlap between each combination of gene pairs (Figure 1C, left). We established that *Nos1*, *Ndnf* and *Gabrr3* cells represent 55%, 70% and 65% of STN^Pitx2^ neurons, respectively. Additionally, more than half of *Nos1 (54%)* and *Ndnf (66%)* positive cells exhibited extensive overlap between them. Finally, a higher percentage of *Gabrr3*-positive cells expressed *Ndnf*+ (86%) than *Nos1+ (59%)* (Figure 1c, right). In sum, (i) *Pitx2* is a general marker for STN neurons, (ii) *Ndnf*- and *Nos1*-expressing neurons represent two partially overlapping clusters of STN neurons and (iii) *Gabrr3*-expressing neurons represent a smaller subpopulation accounting for ~60% of the total population, overlapping with both *Nos1* and *Ndnf* cells (Figure 1C, right; Supplem. Table 2).

### Optogenetic Activation of STN Neurons Decreases Movement of the Contralateral Side

In order to investigate if direct manipulation of STN neurons can regulate motor functions, we conducted neuromodulatory experiments in Pitx2-Cre mice using optogenetics. We injected Cre-dependent channelrhodopsin 2 (AAV5-EF1a-DIO-ChR2-EYFP) in the left STN, to activate STN^Pitx2^ neurons and evaluate their effect on movement (Figure 2A). We adopted the rotation assay, a common test of locomotor function in mice (35,36). Unilateral optogenetic stimulation of STN^Pitx2^ neurons induced motor deficits in the contralateral (right) limbs and thus induced animals to rotate leftward toward the side of the stimulation. Rotations were measured as the ratio between anti-clockwise (leftward) to clockwise (rightward) turns (positive values) and clockwise to anti-clockwise (negative values) (repeated measures two-way ANOVA with Sidak’s multiple comparison test; On epoch: ChR2 mean 29.17 Vs. Control mean −0.71, **** p < 0.0001, n = 6/group; Figure 2C). Interestingly, the observed effect was scalable, with a maximum effect reached at 40 Hz (t-test: increase of mean Control Vs. 10 Hz: 19-fold, * p < 0.05; Vs 40 Hz: mean increse 29-fold, *** p < 0.001; Vs. 120 Hz: 13-fold, ** p < 0.01; n = 6/group; Figure 2B). On the other hand, photoinhibition of STN^Pitx2^ neurons with inhibitory archaerhodopsin (eArch3.0), did not affect the rotation ratio (repeated measures two-way ANOVA with Sidak’s multiple comparison test; Stimulation On epoch: eArch3.0 mean 0.80 Vs. Control mean −0.93, *n.s.* p > 0.05, n = 6/group; Figure 2D). Finally, we tested whether selective ablation of STN^Pitx2^ neurons with diphtheria toxin subunit A (AAV1-EF1a-DIO-dtA) affects rotation behavior. Consistent with photoinhibition, no difference was observed between STN^Pitx2^ ablated mice and control virus injected mice (t-test: dtA mean 1.11 Vs. Control mean −0.70, *n.s.* p > 0.05, n = 6-9/group; Figure 2E). Overall, these data demonstrate that direct activation of the STN through STN^Pitx2^ neurons can regulate motor functions, while acute inhibition or ablation of these neurons is insufficient to induce unilateral rotations.

**Figure 2.**
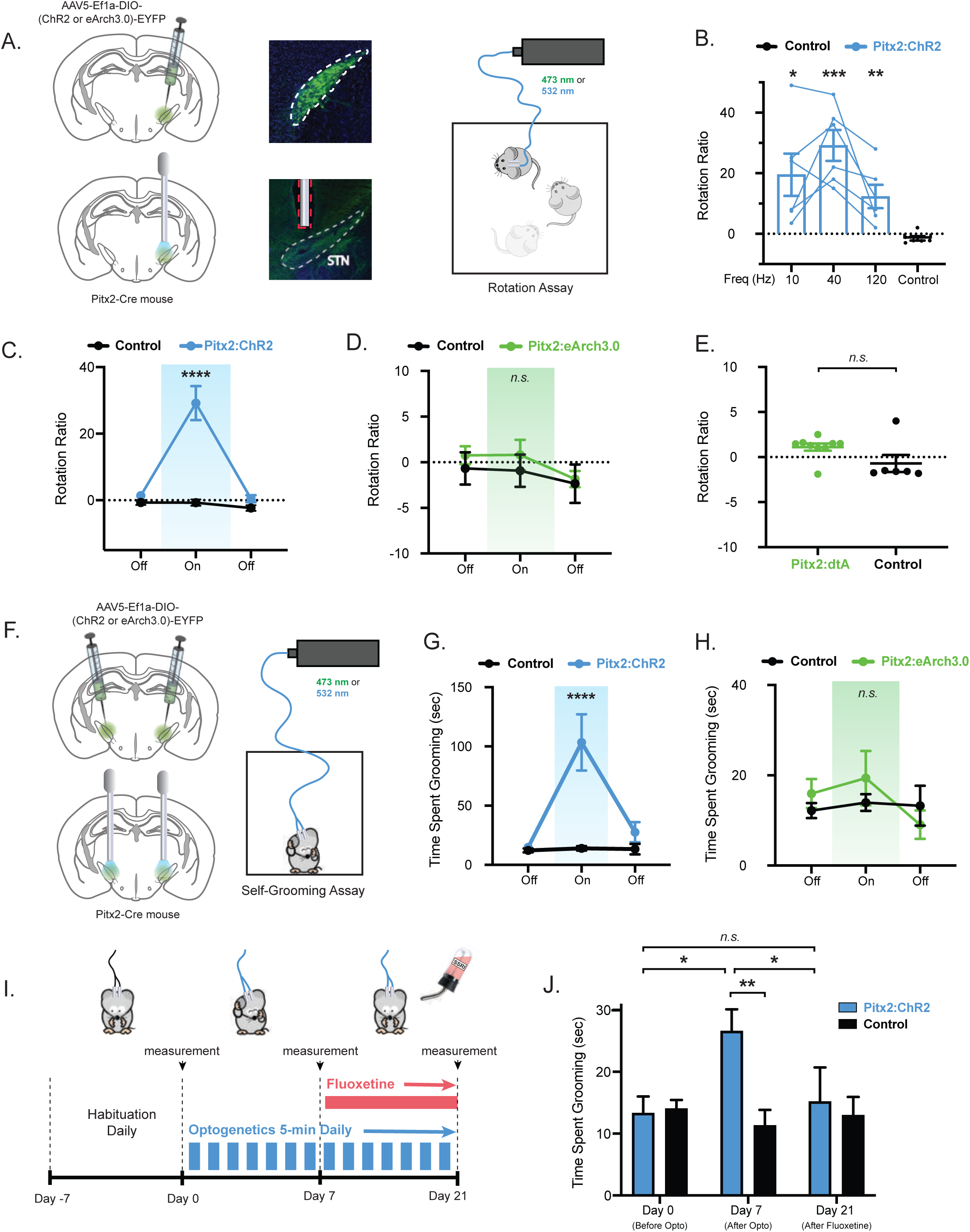
Optogenetic Activation of STN Neurons (STN^Pitx2^) Inhibits Locomotion and Induces OCD-like Repetitive Behavior in Mice. (A) Schema for unilateral optogenetic modulation of STN^Pitx2^ neurons in the Rotation Assay, and representative IHC for validation of viral expression and optic fiber placement. (B) Optogenetic photoactivation of STN^Pitx2^ neurons in freely moving mice significantly increases unilateral rotation ratio. T-student test comparing the effect of stimulation at 10, 40 and 120 Hz to control (* p < 0.05, ** p < 0.01, *** p < 0.001; n = 6 mice per group). (C and D) Photoactivation (40 Hz) of Pitx2:ChR2 mice increases unilateral rotations compared to control mice (C), whereas photoinhibition of Pitx2:eArch3.0 does not alter the rotation ratio compared to control (D). Repeated-measures two-way ANOVA, followed by *ad hoc* Sidak’s multiple comparison test comparing treated and control groups (**** p < 0.0001, *n.s.* p > 0.05; n = 6 mice per group). Blue-shaded region highlights 532nm Laser On epoch. Green-shaded region highlights 473 nm Laser On epoch. (E) Unilateral ablation of STN^Pitx2^ neurons with diphtheria toxin A (dtA) does not alter the rotation ratio of treated mice compared to controls. T-student test comparing Pitx2:dtA to control (*n.s.* p > 0.05; n = 6-9 mice per group). (F) Schema for bilateral optogenetic modulation of STN^Pitx2^ neurons in the Self-Grooming Assay. (G and H) Photoactivation (40 Hz) of Pitx2:ChR2 mice bilaterally increases the time spent grooming acutely (during Laser “On” epoch) compared to control mice (G), whereas bilateral photoinhibition of Pitx2:eArch3.0 does not alter the grooming time compared to control (D). Repeated-measures two-way ANOVA, followed by *ad hoc* Sidak’s multiple comparison test comparing treated and control groups (**** p < 0.0001, *n.s.* p > 0.05; n = 6 mice per group). Blue-shaded region highlights 532nm Laser On epoch. Green-shaded region highlights 473 nm Laser On epoch. (I) Schema for repeated bilateral optogenetic modulation of STN^Pitx2^ neurons in the Self-Grooming Assay. After 7 days of habituation to the optic fiber, Pitx2:ChR2 and control mice were photoactivated (40 Hz) for 5 minutes daily for 7 days. The photostimulation paradigm was then continued for an additional 14 days, while mice were treated with SSRI fluoxetine. Self-grooming time was measured before (Day 0) and after (Day 7) the optogenetic stimulation paradigm, and after the treatment with fluoxetine (Day 21). (J) Photoactivation (40 Hz) of Pitx2:ChR2 mice bilaterally for 7 consecutive days increases the time spent grooming at baseline (24 hours after last Laser On epoch) compared to self at Day 0, and to control mice at Day 7, while the addition of treatment with SSRI fluoxetine reverts the grooming time back to initial levels. Repeated-measures two-way ANOVA, followed by *ad hoc* Sidak’s multiple comparison test comparing treated and control groups, and treated group between Day 0, 7 and 21 (** p < 0.01, * p < 0.05, *n.s.* p > 0.05; n = 6 mice per group). All data are presented as mean ± SEM.

### Acute and Chronic Activation of STN Neurons Elicits Grooming

Extensive evidences have established the STN as a key node in regulating limbic-associative functions and OCD (32). Previous reports have shown that inhibition of the STN with high frequency stimulation suppresses excessive self-grooming in mouse models of autism (21). To evaluate this effect, we initially tested the effect of bilateral photoactivation of STN^Pitx2^ neurons at optimal 40 Hz frequency. Freely moving mice were blindly scored for the time spent grooming during a 15-minute session (5-min pre-stimulation, 5-min stimulation “On”, 5-min post-stimulation) (Figure 2F). Strikingly, a significant increase of highly repetitive self-grooming behavior was observed comparing the stimulation period to either the pre and post stimulation period. Littermate controls did not exhibit such effects in grooming (repeated measures two-way ANOVA with Sidak’s multiple comparison test; Stimulation On epoch: ChR2 mean 103.4s Vs. Control mean 13.96s, **** p < 0.0001, n = 6/group; Figure 2G) (Supplementary Video 1). Inhibition of STN^Pitx2^ neurons with eArch3.0 did not affect grooming behavior between treated and control groups (repeated measures two-way ANOVA with Sidak’s multiple comparison test; Stimulation On epoch: eArch3.0 mean 19.37s Vs. Control mean 13.96s, *n.s.* p > 0.05, n = 6/group; Figure 2H). Importantly, this repetitive behavior was extinguished almost immediately after photostimulation ceased. This suggested that inducing a more sustained pattern of repetitive, stereotyped behavior, independent of acute photostimulation, would replicate OCD features more closely.

Prior studies have shown that optogenetic activation of a cortico-striatal sub-circuit can increase baseline grooming in mice only after several daily repetitions (33). Furthermore, DBS and selective serotonin reuptake inhibitors (SSRI) drug treatments of OCD require days or weeks to become effective (6,7). This suggests that repeated activation could lead to sustained grooming behavior even after photostimuation has ceased. We thus analyzed grooming after repeated photo-stimulations of STN^Pitx2^ neurons over 7 consecutive days and compared the amount of time spent grooming before and after the week-long paradigm. (Figure 2I). We observed a significant increase in grooming during a 5-minute period 24 hours after the final bout of photostimulation (repeated measures two-way ANOVA with Sidak’s multiple comparison test; Day 7 ChR2 Vs. Control: mean increase 2.34-fold, * p < 0.05, ** p < 0.01; ChR2 Day 0 Vs. Day 7: mean increase 2.1 -fold, * p < 0.05, n = 6/group; Figure 2J). Computerized grooming recording over 2 hours confirmed this data in a separate set of experiments (t-test Day 7: ChR2 mean 2350s Vs. Control mean 1679s, * p < 0.05, n = 6/group; Supplementary Figure 1A). Next, to rule out locomotion as a cause for the observed increase in grooming, we quantified the distance traveled by mice and observed no change between treated and control groups (t-test Day 7: ChR2 mean 368 cm Vs. Control mean 333 cm, *n.s.* p > 0.05, n = 5/group; Supplementary Figure 1B). Similar to the effects on rotations, 7 days of repeated photoinhibition of STN^Pitx2^ neurons using eArch3.0 did not alter baseline grooming or total locomotion (repeated measures two-way ANOVA with Sidak’s multiple comparison test, left; t-test, center and right; eArch3.0 Vs. Control, *n.s.* p > 0.05, n = 6/group; Supplementary Figure 1C).

To confirm that the increased grooming represented an OCD-like phenotype, we tested whether drugs used to treat OCD could reverse this photostimulation-induced repetitive behavior. SSRIs like fluoxetine are first-line treatment for OCD patients and have been shown to successfully reduce excessive grooming in mouse models of OCD after two weeks of daily treatment (37,38). Therefore, after inducing over-grooming with 7 successive daily photostimulations of STN^Pitx2^ neurons, we tested whether fluoxetine treatment (20mg/kg) could reverse the behavioral effects of daily stimulation (Figure 2I). Importantly, 14 days of fluoxetine treatment in conjunction with repeated stimulation significantly restored the grooming time to levels comparable to baseline, as analyzed both manually (repeated measures two-way ANOVA with Sidak’s multiple comparison test; Day 21: ChR2 mean 15.17s Vs. Control 12.98s, *n.s.* p > 0.05; ChR2 Day 0 Vs. Day 21, *n.s.* p > 0.05; ChR2 Day 7 Vs. Day 21, 1.8-fold mean decrease, * p < 0.05; n = 6/group; Figure 2J) and with an automated computer system (repeated measures two-way ANOVA with Sidak’s multiple comparison test; ChR2 mean Day 7: 2350s Vs. Day 21: 1621s, ** p < 0.01; n = 6/group Supplementary Figure 1A).

Taken together, these results indicate that STN neurons can be photostimulated to induce an acute increase in repetitive grooming. Moreover, repeated photostimulation induces a persistent over-grooming phenotype, similar to OCD in mice, which can be reversed with fluoxetine treatment.

### Functional Studies of Subpopulations of STN Neurons

Next, to determine whether subpopulations of STN^Pitx2^ neurons (see Figure 1) could recapitulate the observed behavioral responses, we injected a Cre-dependent AAV expressing ChR2 in into the STN of Nos1-Cre, Ndnf-Cre and Gabrr3-Cre mice (Figure 3A). Similar to the results using Pitx2-Cre mice, unilateral photostimulation at 40 Hz of each of these three subpopulations induced a significant change in rotation (t-test: increase of mean Control Vs. Ndnf: 8.6-fold, * p < 0.05; Vs Nos1: 6.9-fold, ** p < 0.01; Vs. Gabrr3: 7.4-fold, ** p < 0.01; n = 4-6/group; Figure 3B). Additionally, except in *Nos1*-Cre mice, we could also recapitulate the acute effects on grooming (t-test: increase of mean Control Vs. Ndnf: 6.7-fold, * p < 0.05; Vs Nos1: 4.9-fold, *n.s.* p > 0.05; Vs. Gabrr3: 7.6-fold, * p < 0.05; n = 4-6/group; Figure 3C) (Supplementary Video 3). Altogether, these data demonstrate that while activation of each of the three subpopulations exhibits qualitatively analogous behaviors, activation of *Gabrr3* and *Ndnf* expressing neurons have stronger effects on rotation and grooming than activation of *Nos1* neurons. Additionally, *Gabrr3* is a smaller cell population and is more specific to the STN than *Ndnf*. Finally, and most importantly, *Gabrr3* is a gene that encodes for a membrane GABA receptor and therefore could be an interesting druggable candidate for translational work.

**Figure 3.**
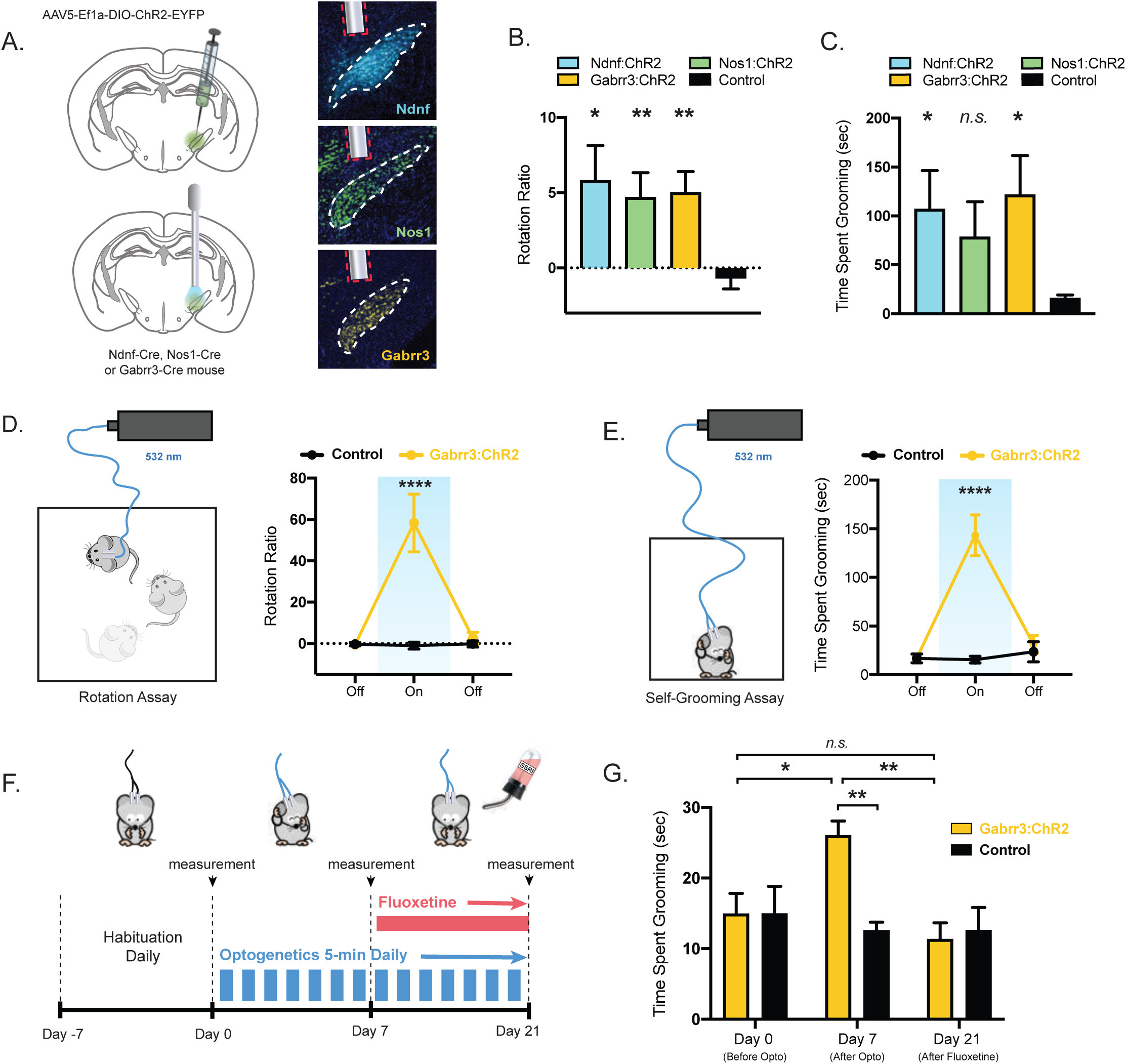
Optogenetic Activation of Gabrr3-Expressing STN Neuronal Subpopulation (STN^Gabrr3^) Inhibits Locomotion and Induces OCD-like Repetitive Behavior in Mice. (A) Schema for unilateral optogenetic modulation of STN neuronal subpopulations in the Rotation Assay, and representative IHC for validation of viral expression and optic fiber placement in Ndnf:ChR2, Nos1:ChR2 and Gabrr3:ChR2 mice. (B and C) Optogenetic photoactivation (40 Hz) of Ndnf, Nos1 and Gabrr3 STN neuronal subpopulations increases unilateral rotations (B), while it significantly increases the time spent grooming only in Ndnf and Gabrr3 populations. T-student test comparing each treated group to control (* p < 0.05, ** p < 0.01, *n.s.* p > 0.05; n = 4-6 mice per group). (D and E) Schema and data from photoactivation (40 Hz) of STN^Gabrr3^ neurons in the Rotation and Self-Grooming Assays; unilateral photoactivation of Gabrr3:ChR2 mice increases unilateral rotations compared to control mice (D), and bilateral photoactivation increases the time spent grooming compared to control mice (D). Repeated-measures two-way ANOVA, followed by *ad hoc* Sidak’s multiple comparison test comparing treated and control groups (**** p < 0.0001; n = 5-6 mice per group). Blue-shaded region highlights 532nm Laser On epoch. (F) Schema for repeated bilateral optogenetic modulation of STN^Gabrr3^ neurons in the Self-Grooming Assay. After 7 days of habituation to the optic fiber, Gabrr3:ChR2 and control mice were photoactivated (40 Hz) for 5 minutes daily for 7 days. The photostimulation paradigm was then continued for an additional 14 days, while mice were treated with SSRI fluoxetine. Self-grooming time was measured before (Day 0) and after (Day 7) the optogenetic stimulation paradigm, and after the treatment with fluoxetine (Day 21). (G) Photoactivation (40 Hz) of Gabrr3:ChR2 mice bilaterally for 7 consecutive days increases the time spent grooming at baseline (24 hours after last Laser On epoch) compared to self at Day 0, and to control mice at Day 7, while the addition of treatment with SSRI fluoxetine reverts the grooming time back to initial levels. Mixed-model two-way ANOVA, followed by *ad hoc* Sidak’s multiple comparison test comparing treated and control groups, and within groups between Day 0, 7 and 21 (* p < 0.05, ** p < 0.01, *n.s.* p > 0.05; n = 5-6 mice per group). All data are presented as mean ± SEM.

We thus decided to evaluate the effect of modulating Gabrr3-expressing neurons (hereafter STN^Gabrr3^) activity in finer detail. First, we confirmed that 40 Hz is the optimal frequency for STN^Gabrr3^ photostimulation (t-test: increase of mean Control Vs. 10 Hz: 27-fold, * p < 0.05; Vs 40 Hz: mean increase 68-fold, *** p < 0.001; Vs. 120 Hz: 17-fold, *n.s.* p > 0.05; n = 5-6/group; Supplementary Figure 2A). Importantly, unilateral photostimulation of STN^Gabbr3^ neurons at this frequency acutely induced a significant increase in ipsilateral rotations. These rotations were quantitatively double to the response to STN^Pitx2^ photoactivation (repeated measures two-way ANOVA with Sidak’s multiple comparison test; On epoch: ChR2 mean 58.33 Vs. Control mean −1.23, **** p < 0.0001, n = 5-6/group; Pitx2 29.17 Vs. Gabrr3 58.33; Figure 3D). Similarly, bilateral photostimulation significantly induced an increase in grooming. This effect was also stronger than when evaluating Pitx2 activation (repeated measures two-way ANOVA with Sidak’s multiple comparison test; On epoch: ChR2 mean 143.3 s Vs. Control mean 15.44, **** p < 0.0001, n = 5-6/group; Pitx2 103.4s Vs. Gabrr3 143.3s; Figure 3E). These results further support *Gabrr3* as a particularly valuable marker for STN neuronal modulation of both PD- and OCD-like behaviors.

To determine whether the chronic stimulation paradigm led to spontaneous and persistent grooming even after photo-stimulation had ceased also in the STN^Gabrr3^ subpopulation (Figure 3F), we repeated photostimulation of these neurons for 7 days and evaluated grooming. Indeed, we observed a significant increase in grooming time in the treated group at day 7 (mixed-effect analysis with Sidak’s multiple comparison test; Day 7 ChR2 Vs. Control: mean increase 2.1-fold, ** p < 0.01; ChR2 Day 0 Vs. Day 7: mean increase 1.8-fold, * p < 0.05, n = 5-6/group; Figure 3G). These data were confirmed using an automatized computer system (t-test Day 7: ChR2 mean 2216s Vs. Control mean 1539s, * p < 0.05, n = 5-6/group; Supplementary Figure 2B). Similar to STN^Pitx2^ neurons repeated stimulation of STN^Gabbr3^ did not alter total locomotion (t-test Day 7: ChR2 mean 1685 cm Vs. Control mean 1535 cm, *n.s.* p = 0.67, n = 5-6/group; Supplementary Figure 2C). As expected, treatment with fluoxetine for two weeks suppressed the repetitive grooming behavior as measured both manually (mixed-effect analysis with Sidak’s multiple comparison test; Day 21: ChR2 mean 11.33s Vs. Control 12.61s, *n.s.* p = 0.98; ChR2 Day 7 Vs. Day 21: mean decrease 2.3-fold, ** p < 0.01, n = 5-6/group; Figure 3G), and with the automated computerized system (t-test Day 21: ChR2 mean 1575s Vs. Control mean 1597s, *n.s* p = 0.97, n = 5-6/group; Supplementary Figure 2B).

Taken together, these results suggest that modulation of the activity of STN^Gabrr3^ neurons recapitulates the behavior observed when manipulating the entire STN (Pitx2), but has a quantitatively stronger effect.

### Modulation of STN^Gabrr3^ Neurons in Mouse Models of Parkinson’s Disease and OCD

Our results showing that activation of STN^Gabrr3^ neurons could lead to rotations and induce repetitive grooming behavior suggested that modulating the activity of these neurons might improve the phenotypes of common mouse models of PD and OCD. To test this possibility in PD models, we first modulated STN^Gabrr3^ neuronal activity in mice rendered hemi-parkinsonian by unilateral injection of the toxic dopamine analogue 6-hydroxydopamine (6-OHDA) into the left medial forebrain bundle (MFB) (60). When injected in the MFB, 6-OHDA induces dopaminergic neuronal ablation resulting in a drastic decrease of dopamine marker tryptophan hydroxylase (TH) in the striatum of the injected side (Figure 4A). As a consequence, mice display deficits in the contralateral (right) side of the body and rotate in a direction ipsilateral to the lesion (leftward). Moreover, administration of amphetamine, by increasing the synaptic release of dopamine, exacerbates the effects and facilitates functional evaluation.

**Figure 4.**
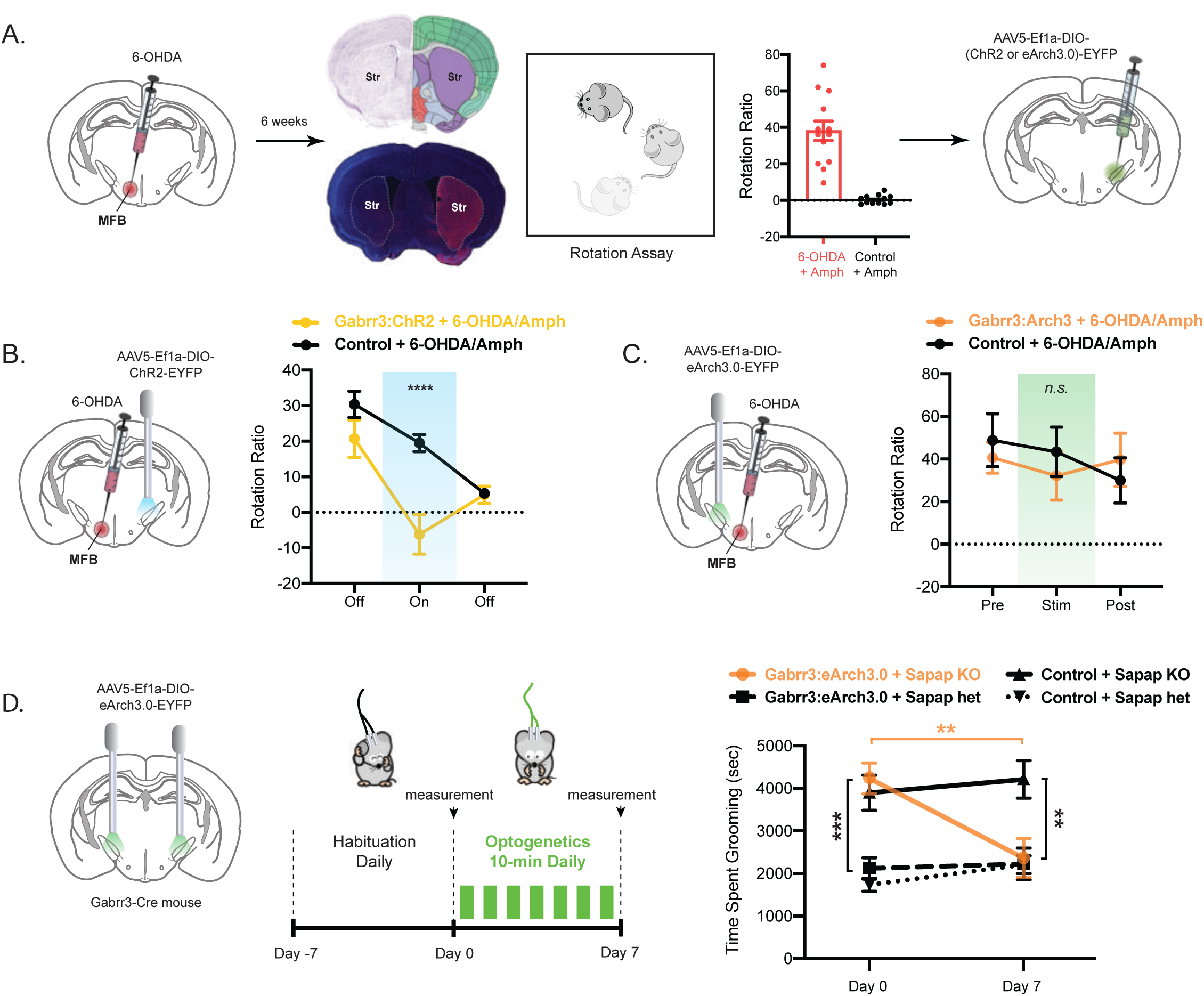
Optogenetic Modulation of STN^Gabrr3^ Neurons Improves the Pathologic Phenotypes of PD and OCD Mouse Models. (A) Schema of induction of hemi-parkinsonian phenotype with 6-OHDA. Injection of 6-OHDA in the left medial forebrain bundle (left) induces ipsilateral dopaminergic neuronal ablation resulting in a drastic decrease of dopamine marker TH in the striatum; this induces an increase in ipsilateral rotations compared to controls, which is exacerbated by amphetamine (Amph) (center). Individual animals scoring Rotation Ratio > 5 were selected to undergo optogenetics surgery (right). (B and C) Schema and data showing that photoactivation of STN^Gabrr3^ neurons contralateral to the 6-OHDA lesion rescues the hemi-parkinsonian rotation behavior of the treated group to normal levels (B), whereas photoinhibition (40 Hz) of ipsilateral STN^Gabrr3^ neurons did not change the rotation behavior of hemi-parkinsonian mice compared to controls (C). Repeated-measures two-way ANOVA, followed by *ad hoc* Sidak’s multiple comparison test comparing treated and control groups (**** p < 0.0001, *n.s.* p > 0.05; n = 5-8 mice per group). Green-shaded region highlights 473 nm Laser On epoch. Blue-shaded region highlights 532nm Laser On epoch. (D) Schema and data for repeated bilateral optogenetic inhibition of STN^Gabrr3^ neurons in Sapap3-KO mice in the Self-Grooming Assay. After 7 days of habituation to the optic fiber, Gabrr3:ChR2 and control mice of both Sapap3 KO and Sapap3 heterozygous genotype were photoinhibited for 10 minutes daily for 7 days. Self-grooming time was significantly decreased in the Sapap3 KO treated mice after photoinhibition, while it was not significantly altered in the three control groups (Sapap3 KO control, Sapap3 heterozygous treated, Sapap3 heterozygous control). Repeated-measures two-way ANOVA, followed by *ad hoc* Tukey’s multiple comparison test comparing treated and control groups, and comparing before and after photostimulation within each group (** p < 0.01, *** p < 0.001, n.s. non-significant; n = 5-7 mice per group). All data are presented as mean ± SEM.

Our results show that in 6-OHDA-amphetamine-treated mice photostimulation of contralateral STN^Gabrr3^ neurons, opposite to the side of the lesion, restored the ipsilateral rotations to normal levels (repeated measures two-way ANOVA with Sidak’s multiple comparison test; Stimulation On epoch: ChR2 mean −6.21 Vs. Control mean 19.50, **** p < 0.0001, n = 8/group Figure 4B). Similar to the lack of effect when inhibiting Pitx2 neurons, photoinhibition of STN^Gabrr3^ neurons ipsilateral to the lesion failed to diminish the leftward-to-rightward rotation ratio to baseline in amphetamine-treated animals (repeated measures two-way ANOVA with Sidak’s multiple comparison test; Stimulation On epoch: eArch3.0 mean 32.16 Vs. Control mean 43.40, *n.s* p > 0.05, n = 5/group Figure 4C).

Moreover, in order to evaluate a role for STN^Gabrr3^ neurons in OCD, we tested the effect of repeated photomodulation of STN^Gabrr3^ neurons on Sapap3 KO mice, an established animal model of OCD (38,39). Because repeated photoactivation of STN^Gabrr3^ neurons increases baseline grooming (Figure 3G), we hypothesized that repeated photoinhibition of the same neurons could decrease the pathological over-grooming in Sapap3 KO mice (Figure 4D, left). Indeed, bilateral photoinhibition of STN^Gabrr3^ for 10-min/day over 7 days significantly decreased the time Sapap3 KO mice spent grooming to the same level as their phenotypically normal, heterozygous littermates (repeated measures two-way ANOVA with Sidak’s multiple comparison test; eArch3.0 mean Day 0: 4330s Vs. Day 7: 2449s, ** p < 0.01, n = 5-9/group; Figure 4D, right). These data demonstrate that modulation of STN^Gabrr3^ neuronal activity can ameliorate the symptoms associated with PD and OCD, and highlights *Gabrr3* in the STN as an interesting marker for neuronal modulation and drug development.

### Anatomic and Functional Mapping of STN^Gabrr3^ Projections

We next set out to identify the targets of STN^Gabrr3^ to evaluate whether this population of neurons is projecting to the known targets of the STN. To do so, we injected a Cre-dependent AAV expressing the red fluorescent protein mCherry fused to a transmembrane receptor (AAV-hSyn1-DIO-hM3(Gq)-mCherry) into the STN of Gabrr3-Cre mice (Figure 5A, top). Six weeks after the injection, which enabled expression and transport of mCherry through neuronal axons, we used the whole-brain immunolabeling approach iDISCO+ paired to tissue clearing (29), enabling us to visualize both STN^Gabrr3^ soma and axon terminals in optical sections of whole brains (Figure 5A, bottom left). Importantly, we found that dense projections of STN^Gabrr3^ neurons where found both in the entopeduncular nucleus (EP) and globus pallidus (GP) anteriorly, and to the substantia nigra (SNr) posteriorly (Figure 5A, center and right). This projection pattern is similar to that of the STN as a whole (40). We next asked whether the STN^Gabrr3^ neurons that project to the SNr and GP/EP represent distinct, partially overlapping, or the same cell populations. To address this, we performed a dual color tracing approach using retrogradely transported AAVs expressing different Cre-dependent fluorescent proteins. We used Gabrr3-Cre mice and injected an AAV expressing a Cre-dependent green fluorescent protein (GFP) into the GP/EP, and a second AAV expressing a Cre-dependent red fluorescent protein (RFP) into the SNr. As a result, STN^Gabrr3^ neurons projecting to the GP/EP express GFP, STN^Gabrr3^ neurons projecting to the SNr express RFP, and neurons projecting to both sites are yellow (Figure 5B). Quantitative analysis of the images revealed that 27% of STN^Gabrr3^ neurons project exclusively to the GP/EP (green), 37% of STN^Gabrr3^ project to the SNr (red) and 36% project to both sites (yellow) (Figure 5C). These data suggest that STN^Gabrr3^ neurons project equally to the two known targets of the STN, and that more than one third of the population sends collateral projections to both.

**Figure 5.**
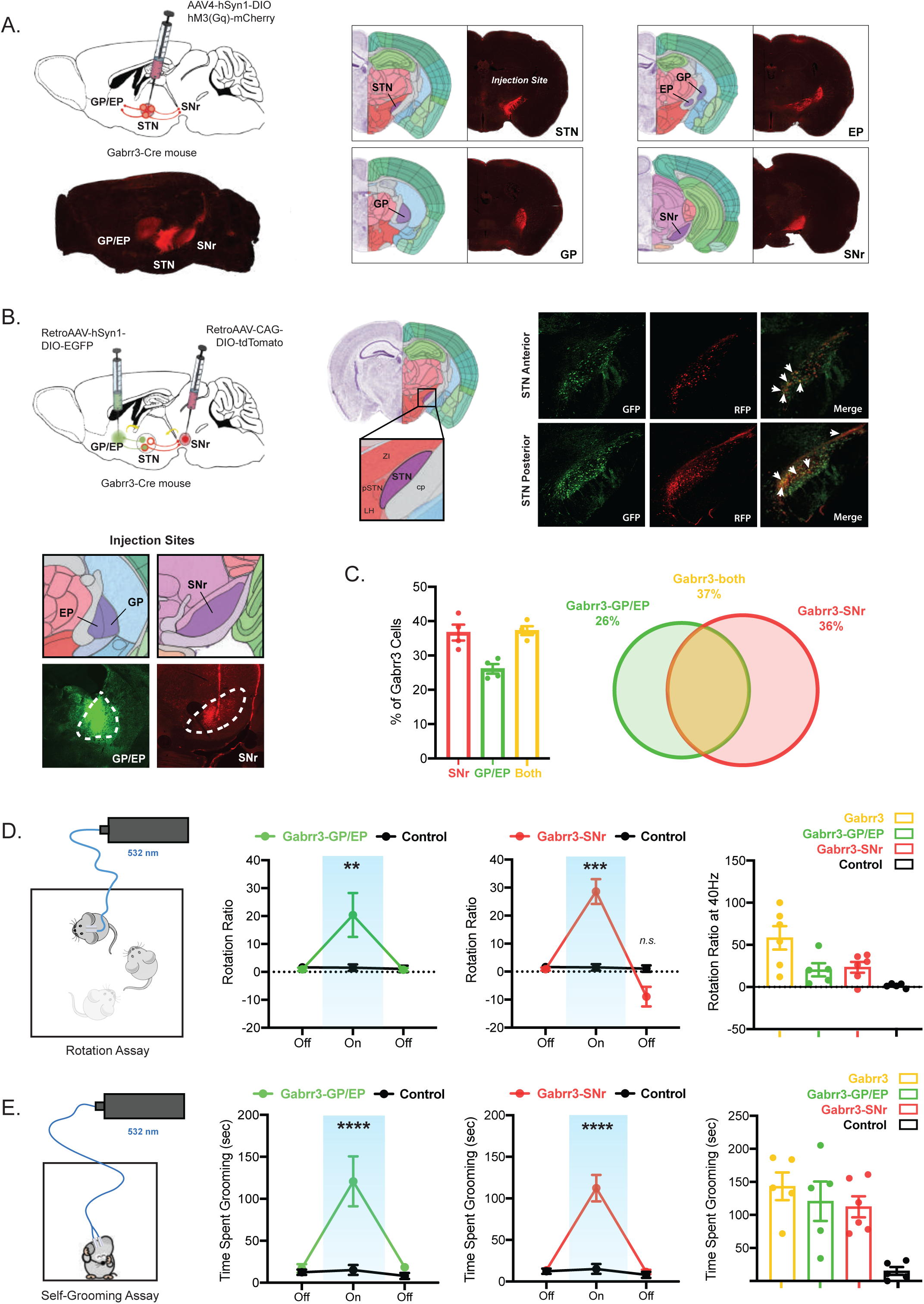
STN^Gabrr3^ Neurons Regulate Locomotion and Repetitive Behavior Via Projections to the GP/EP and SNr. (A) Schema and representative images of whole-brain projection mapping of STN^Gabrr3^ neurons. Left, top: projections were traced via injection of a Cre-dependent AAV expressing a red fluorescent protein in the STN of Gabrr3-Cre mice. Left, bottom: optical sagittal section of whole-brain cleared with iDISCO+ showing injection site (STN) and projections (GP/EP, SNr.) Center and right: left panels represent Allan Brain Atlas annotations for STN, EP, GP and SNr; right panel represent coronal sections from iDISCO+ experiment of the respective brain areas. (B) Schema and representative images of dual color retrograde tracing from STN^Gabrr3^ neuronal terminals. Left, top: neuronal terminals were infected with Cre-dependent AAVs expressing RFP in the SNr, and GFP in the GP/EP of Gabrr3-Cre mice. Left, bottom: representative IHC and respective Allan Brain Atlas annotations confirming anatomically accurate surgical targeting of terminals. Right: representative IHC images of the STN showing STN^Gabrr3^ neurons labeled in green, red or both (yellow, white arrows) in the anterior and posterior parts of the STN. (C) Quantification of percentage of STN^Gabrr3^ neurons projecting to the GP/EP (green), SNr (red) and both (yellow), and Venn diagram summarizing quantitative overlapping of the neuronal sub-populations based on projections. (D and E) Schema and data from photoactivation (40 Hz) of STN^Gabrr3^ projections in the Rotation and Self-Grooming Assays. (D) Unilateral photoactivation of Gabrr3:ChR2 projections to GP/EP (center-left panel) and SNr (center-right panel) increases rotations compared to control. Right panel: compared effect between groups during Laser On epoch. (E) Bilateral photoactivation of STN^Gabrr3^ neurons projecting to GP/EP (center-left panel) or SNr (center-right panel) increase the time spent grooming compared to control. Right panel: compared effect between groups during Laser On epoch. Repeated-measures two-way ANOVA, followed by *ad hoc* Sidak’s multiple comparison test comparing treated and control groups (** p < 0.01, *** p < 0.001, **** p < 0.0001, n.s. non-significant; n = 5-6 mice per group). Blue-shaded region highlights 532nm Laser On epoch. All data are presented as mean ± SEM.

Finally, we tested whether STN^Gabrr3^ projections are functionally distinct. To do so, we injected a Cre-dependent AAV expressing ChR2 in the STN of Gabrr3-Cre mice, and placed an optic fiber at each of the projection sites (GP/EP or SNr). We analyzed the number of rotations in response to unilateral photostimulation, and the grooming behavioral assay during bilateral photostimulation at each of these sites. We found that photoactivation of projections to both the GP/EP and the SNr significantly increased the number of unilateral rotations (repeated measures two-way ANOVA with Sidak’s multiple comparison test; On epoch mean: Gabrr3-SNr 23.33 Vs. Control 1.533 *** p < 0.001, n = 5-6/group; Gabrr3-GP/EP 20.40 Vs. Control 1.533 ** p < 0.01, n = 5/group; Figure 5D). However, the magnitude of the effect was significantly smaller than the photoactivation of the entire STN^Gabrr3^ population (one-way ANOVA with multiple comparisons: On epoch mean: Gabrr3 58.33 Vs. Gabrr3-SNr 23.33, * p < 0.05; Gabrr3 58.33 Vs. Gabrr3-GP/EP 20.40, * p < 0.05; n = 5-6/group; Figure 5D, right). Similarly, photostimulation at each of these sites increased grooming (repeated measures two-way ANOVA with Sidak’s multiple comparison test; On epoch mean: Gabrr3-SNr 112.3s Vs. Control 15.15s **** p < 0.0001, n = 5-6/group; Gabrr3-GP/EP 120.80s Vs. Control 15.15s **** p < 0.0001, n = 5/group; Figure 5E). This increase was less than that induced by photoactivation of the entire STN^Gabrr3^ population, but not significantly smaller (one-way ANOVA with multiple comparisons: mean On epoch: Gabrr3 143.3s Vs. Gabrr3-SNr 112.3s, *n.s.* p = 0.71; Gabrr3 143.3s Vs. Gabrr3-GP/EP 120.80s, *n.s.* p = 0.51; n = 5-6/group). Importantly, the magnitude of effect was similar after activation of STN^Gabrr3^ terminals in the GP/EP and SNr projections, further suggesting that both projections can regulate locomotor and limbic-associative functions (t-test Gabrr3-SNr Vs. Gabrr3-GP/EP: rotations, *n.s.* p > 0.05; grooming, *n.s.* p > 0.05; Figure 5D and E, right). In aggregate, these results suggest that STN^Gabrr3^ neurons mediate their limbic-associative and motor functions through projections to both the GP/EP and SNr, and they do not present a projection-dependent functional segregation.

## DISCUSSION

Since its introduction in the clinic twenty years ago, over 160 000 patients with PD and other neuropsychiatric disorders have benefitted from DBS treatment. The most common target of this procedure in PD is the STN (4,5). Furthermore, DBS-STN can also provide supplemental relief in patients with OCD (41). While the available evidence suggests that high frequency DBS acts by inactivating the STN, this procedure can also alter the activity of brain areas adjacent to the STN and/or local fibers of passage (25,42). This occurs because DBS is not cell-specific, thus the correct surgical placement of electrodes contributes greatly to the variability in efficacy and side effects observed between patients (43). It is therefore of basic and potential clinical importance to define the neuronal populations responsible for DBS’s effects, which could enable the development of alternative cell-specific therapeutic approaches.

Although the STN has been widely studied in health and disease, its cell populations and their role on PD pathogenesis are poorly understood. Our data (Figure 1), in addition to published work (34,44), demonstrate that most STN neurons are glutamatergic and express the transcription factor *Pitx2*. Prior to this report, specific STN neuronal subpopulations have not been characterized. Here we identify three overlapping sets of STN neurons expressing *Ndnf*, *Nos1* and *Gabrr3*. Modulation of these neuronal clusters improves PD- and OCD-like phenotypes in mice, with the stronger effect observed in STN^Gabrr3^ neurons. *Gabrr3* gene encodes for a subunit of the GABA-Aρ receptor, a rare type of GABA-A receptor, and could thus represent a potential target for new pharmacologic treatments.

Several lines of evidence suggest that the STN inhibits movement initiation, and that when it is disinhibited in PD, the resulting hyperactivity is associated with hypokinesis of movements (10,11). It has been further suggested that the symptomatic improvement conferred by DBS in PD patients is the result of STN inhibition (12). The experimental support for this possibility includes the following: i. anatomical lesions of the STN improve hypokinesa (13), ii. DBS-STN in mice decreases the firing rate of STN neurons (45) as well as its excitatory output to the SNr (46), iii. injection of muscimol - a GABA agonist and neuronal inhibitor - in the STN of monkeys increases contralateral limb movements (20), and iv. decreased glutamatergic signaling in STN neurons increases locomotion in mice (34). However, cell-type specific modulation of STN neurons on locomotion or grooming has not been evaluated in mice. Importantly, assessment of unilateral optogenetic manipulation of the STN on rotation in rats did not show an effect either at “low-frequency” (20 Hz) or “high frequency” (120 Hz) photostimulation (35). Because the effect of DBS is frequency-dependent, we asked whether rotation required frequencies different than those that were tested. STN neurons have been reported firing at spontaneous spike frequencies between 5-40 Hz *ex vivo* (47), with an average in rodents of around 10-12 Hz *in vivo* (45). We thus decided to empirically test 10 Hz, as an average low frequency; 40 Hz, the upper limit of spontaneous STN frequency range (48), and 120 Hz, to mimic the frequency used in DBS (35). Intriguingly, our data demonstrate a maximal effect at 40 Hz in both STN^Pitx2^ (Figure 2B) and STN^Gabrr3^ neurons (Supplementary Figure 2A). This suggests that increasing the firing of STN neurons within the physiological range has stronger functional effects than stimulation at higher, non-physiologic frequencies such as those employed in DBS, which may induce detrimental effects including conduction block, synaptic fatigue and antidromic stimulation of afferents (35,42).

Consistent with a previous report (35) acute unilateral photoinhibition of STN^Pitx2^ and STN^Gabrr3^ neurons did not induce unilateral rotations in healthy (Figure 2D) or hemiparkinsonian animals (Figure 4C). It is plausible that STN neurons, while not silent at baseline (45,47), may not exert behavioral effects significant enough to induce rotations. In line with this hypothesis, unilateral ablation of STN neurons in animals using diphtheria toxin also did not induce unilateral rotations (Figure 2E). A second possibility is that, at least acutely, ionotropic inhibition of STN neural activity using optogenetics may not be sufficient to tip the electrophysiological balance between direct and indirect pathways within the basal ganglia neural circuit. In the future, it will thus be interesting to evaluate the effect of other silencing approaches, such as metabotropic inhibition with chemogenetics.

While it is well established that loss of dopaminergic innervation in PD leads to an increase in STN activity (10,11), the changes in dopaminergic innervation and circuit alterations in OCD are less clear. Several lines of evidence point towards an increase in the activity of the cortico-striato-thalamo-cortical circuit (49), and the STN. In fact, electrophysiological studies in OCD patients show a correlation between severity of symptoms, responsiveness to DBS, and overall activity of STN neurons (eg, increased burst frequency and theta oscillatory activity) (50,51). In mice, the Sapap3-KO model of OCD shows defects in excitability more prominent in striato-nigral (direct pathway) than in striato-pallidal neurons (indirect pathway) (52). This suggests that a relative increase in the activity of the indirect pathway activity, and consequently of the STN, may contribute to the symptoms. Consistent with this, high-frequency stimulation of the STN has been proven effective in suppressing self-grooming in mouse models of autism (21) and reduce compulsive lever pressing in rats (22). In line with these extensive evidences, selective and direct optogenetic activation of all STN (Pitx2) neurons or the Gabrr3-expressing subpopulation can induce excessive grooming in mice (Figures 2G, 2J and Figures 3E, 3G), and repeated inhibition of STN^Gabrr3^ cells reduces the over-grooming of Sapap3-KO mice (Figure 4D). Taken together, these data show that direct neuronal modulation of the STN can regulate grooming behavior in mice, and that cell-specific inhibition can be a potential therapeutic approach to treat OCD.

A previous report showed that repeated optogenetic activation of a cortico-striatal sub-circuit increased repetitive grooming in mice even after the activation had stopped (37). In line with this, we observed that repeated photoactivation of STN^Pitx2^ and STN^Gabrr3^ neurons induced sustained grooming behavior even after photostimuation had ceased (Figure 3G). Similarly, in Sapap3-KO mice, repeated photoinhibition of STN^Gabrr3^ neurons decreased pathologic grooming (Figure 4D). These protracted effects are consistent with: (i) inhibition of the STN with DBS suppresses excessive self-grooming in autism-like mice progressively over the 3-5 days post stimulation (21); (ii) chronic but not acute DBS-STN restores cortico-striatal synaptic plasticity and normalizes glutamate and GABA levels in other basal ganglia nuclei, including striatum and SNr (53). Similarly, in the clinic, repeated external modulation is required to achieve therapeutic efficacy in several pharmacologic (eg, SSRI) and neuro-modulatory treatments of psychiatric patients (eg, DBS, electroconvulsive therapy, transcranial magnetic stimulation) (54). Taken together, these and our results suggest that immediate modifications in electrical activity may not be sufficient to correct the aberrant neural activity observed in OCD patients, that instead depends on slower-acting mechanisms, such as changes in neural plasticity and gene expression in the STN and downstream basal ganglia nuclei.

To address this possibility, we analyzed the role of these other basal ganglia circuit elements by mapping the projections of STN^Gabrr3^ populations and by testing their functions. The STN receives dense GABAergic innervation from the GP (indirect pathway) and glutamatergic from the cortex (hyper-direct pathway), and sends glutamatergic projections to the EP in rodents and SNr (10,11,31). Functional and anatomical studies in humans and non-human primates suggest that the STN is divided into three anatomically segregated areas with different functions: limbic, associative and motor (15,20,55). Anatomic studies have suggested that the rodent STN is similarly organized, with a dorsal part modulating motor functions via a GP/EP projection, and a ventral-medial area (corresponding to the paraSTN) regulating limbic-associative functions via projections to the SNr and ventral pallidum (41,56,57). In contrast, other reported that neurons of the paraSTN are involved in limbic functions through projections to the GP (27), not the SNr. To investigate possible anatomic and functional segregation in mice, we first tested whether STN^Gabrr3^ neurons projecting to the GP/EP Vs. SNr are located in anatomically segregated areas of the STN, and if selective photostimulation of different projections induces distinct behaviors. Dual color retrograde tracing from STN^Gabrr3^ terminals in GP/EP and SNr failed to show anatomic segregation, in that neurons of the ventral-medial or dorsal STN did not project preferentially to one target or the other (Figure 5B). Consistent with this, photoactivation of STN^Gabrr3^ projections to the GP/EP or SNr had quantitatively similar effects in inducing rotation and grooming behavior in mice, although the overall effect was lower than activating STN^Gabrr3^ neurons as a whole (Figure 5D and 5E, right). These observations suggest that STN^Gabrr3^ neurons mediate both their motor and limbic-associative functions via projections to the GP/EP and SNr, but also that both projections are needed to convey the full behavioral effect.

In summary, in this study we identified three partially overlapping subpopulations of neurons in the STN and found that optogenetic activation of each induces locomotor deficits and repetitive grooming, both of which are features of PD and OCD. These effects were mediated through projections to the GP/EP and SNr. Optogenetic modulation of STN^Gabrr3^ neurons rescued the behavioral phenotypes of PD and OCD mouse models, consistent with the possibility that DBS acts to improve symptoms in these disorders by directly modulating STN neurons, as opposed to fibers of passage or neighboring areas. Furthermore, and most importantly, these findings also identify *Gabrr3* as a potential therapeutic target in the STN for treating PD and OCD, and may provide a basis for the development of cell-specific therapeutic approaches using chemogenetics or related technologies.

## ACKNOWLEDGMENTS

We thank Dr. Timothy Cox of the University of Washington for contributing the Pitx2-Cre mice, Dr. Marc Flajolet and the Greengard Laboratory at The Rockefeller University for allowing the use of the Home Cage Environment (Clever Sys Inc.), Drs. Claire Henchcliffe, Andrea Lee and the PD & Movement Disorders Institute at Weill Cornell Medicine for consulting on the behavioral phenotype, Drs. Alejandro Lopez and Jessica Jimenez for helpful discussions and manuscript revision, James Knox and Kyle Pellegrino for technical assistance, Anoj Ilanges and Dr. Virginia A. Pedicord for the helpful discussions, and The Rockefeller University Comparative Bioscience Center and Bio-Imaging Resource Center. L.P. acknowledges support from the The David Rockefeller Fellowship and Boehringer Ingelheim Fonds PhD Fellowship. J.M.F acknowledges support from JPB foundation. M.S. acknowledges support from the Kavli NSI Fellowship.

## CONFLICT OF INTEREST

The Authors decleare no conflict of interest.

## FIGURE LEGENDS

**Supplementary Figure 1. Optogenetic Activation of STN Neurons (STN^Pitx2^) Inhibits Locomotion and Induces OCD-like Repetitive Behavior in Mice.**

(A) Optogenetic photoactivation (40 Hz) of Pitx2:ChR2 mice bilaterally for 7 consecutive days increases the time spent grooming at baseline compared to control as measured by automatized computer system (Day 7), while the addition of treatment with SSRI fluoxetine reverts the grooming time back to initial levels (Day 21). Repeated-measures two-way ANOVA, followed by *ad hoc* Sidak’s multiple comparison test comparing treated group between Day 0, 7 and 21 (** p < 0.01, n.s. non-significant; n = 6 mice per group).

(B) Photoactivation (40 Hz) of Pitx2:ChR2 for 7 consecutive days does not change the total locomotion of mice at, measured as total distance walked in 5 minutes. T-student test comparing treated and control groups (n.s. non-significant; n = 6 mice per group).

(C) Photoinhibition of Pitx2:eArch3.0 mice bilaterally for 7 consecutive days does not change the time spent grooming at baseline compared to control as measured manually (left) and by automatized computer system (center), nor the total locomotion over 5 minutes (right). Repeated-measures two-way ANOVA, followed by *ad hoc* Sidak’s multiple comparison test (left), and t-student test comparing treated and control groups (center and right) (n.s. non-significant; n = 6 mice per group).

All data are presented as mean ± SEM.

**Supplementary Figure 2. Optogenetic Activation of Gabrr3-Expressing STN Neuronal Subpopulation (STN^Gabrr3^) Inhibits Locomotion and Induces OCD-like Repetitive Behavior in Mice.**

(A) Optogenetic photoactivation of STN^Gabrr3^ neurons in freely moving mice significantly increases unilateral rotation ratio. T-student test comparing the effect of stimulation at 10, 40 and 120 Hz to control (* p < 0.05, *** p < 0.001; n = 5-6 mice per group).

(B) Photoactivation (40 Hz) of Pitx2:Gabrr3 mice bilaterally for 7 consecutive days increases the time spent grooming at baseline compared to control as measured by automatized computer system (Day 7), while the addition of treatment with SSRI fluoxetine reverts the grooming time back to initial levels (Day 21). T-student test comparing treated and control groups (** p < 0.01, n.s. non-significant; n = 5-6 mice per group).

(C) Photoactivation (40 Hz) of Pitx2:ChR2 for 7 consecutive days does not change the total locomotion of mice, measured as total distance walked in 5 minutes. T-student test comparing treated and control groups (n.s. non-significant; n = 5-6 mice per group).

All data are presented as mean ± SEM.

## Notes

### Competing Interest Statement

The authors have declared no competing interest.

