## Supplemental Tables, Figures for "Functional Analysis of Distinct Populations of Subthalamic Nucleus Neurons on Parkinson’s Disease and OCD-like Behaviors in Mice": 4. Supplementary Table 1-2.pdf (from another version of this preprint: version 1, 2020-06-11)

**Assessment of STN-Specificity of Candidate Marker Genes**

| Gene | Strength | Density (% cells) | Local Sp. | Global Sp. | Overall | Ranking | Notes |
|---|---|---|---|---|---|---|---|
| Pitx2 | +++ | 60-70%; even | +++ | +++ | 9 | 1 | Posterior: transitions to SN, VTA |
| Gabrr3 | +++ | 60-70%; even | +++ | +++ | 9 | 2 | Posterior: transitions to SN, VTA |
| Ndnf | +++ | 60-70%; postero-ventral | ++ | ++ | 9 | 2 | Posterior: transitions to SN |
| Nos1 | ++ | 50% post; 20% antero-dors | ++ | ++ | 9 | 2 | Post: SN sparse; Med: LH lots |
| Nov / CCN3 | +++ | 70% postero-medio-ventral | ++ | ++ | 8 | 3 | Medial: few cells in LH; Dorsal: lots in ctx |
| C130021I20Rik | ++ | 60% even | ++ | +++ | 8 | 3 | Posterior: transitions to SN, VTA |
| Col24a1 | ++ | 30-40%; postero-ventral | ++ | +++ | 7 | 4 | Dorsal: a few + cells in thalamus |
| Kcng4 | ++ | 60%; ventral | ++ | ++ | 7 | 4 | Dorsal/Posterior: very strong in ZI |
| Rapgef5 | ++ | 60-70% even | ++ | ++ | 7 | 4 | Ventral, dorsal: sparse background + ctx |
| Sema3d | +/++ | 30%; medio-ventral | +++ | ++ | 7 | 4 | Weak signal/questionable background |
| Ttc6 | + | 40-50%; even | +++ | ++ | 6 | 5 | |
| Sema3a | +/++ | 20-30%; medio-ventral | ++ | + | 5 | 6 | Dorsal in ctx, poterior very light in SN |
| Ankrd34b | ++ | 50% even | + | + | 5 | 6 | Ventral: ZI, thalamus; Posterior: SN |
| Kcnab3 | ++ | 70% even | + | + | 5 | 6 | Dorsal: ZI + thalamus + ctx |
| Sema3f | + | 30-40%; even | ++ | ++ | 5 | 6 | Medial, dorsal: ZI, LH, thalamus |
| Zfp618 | + | 30%; medio-ventral | ++ | + | 4 | 7 | |
| Stard5 | + | 60-70%; even | + | + | 4 | 6 | High background all around |
| Alox8 | + | 30-40% even | + | + | 3 | 8 | Heavy background |
| Kit / SCFR | +/++ | 30-40% even | + | + | 3 | 8 | Sparse background; heavy hippocampus |
