## Supplementary figures and images for "Functional Analysis of Distinct Populations of Subthalamic Nucleus Neurons on Parkinson’s Disease and OCD-like Behaviors in Mice" (from another version of this preprint: version 1, 2020-06-11)

### 5. Supplementary Figure 1-2.pdf

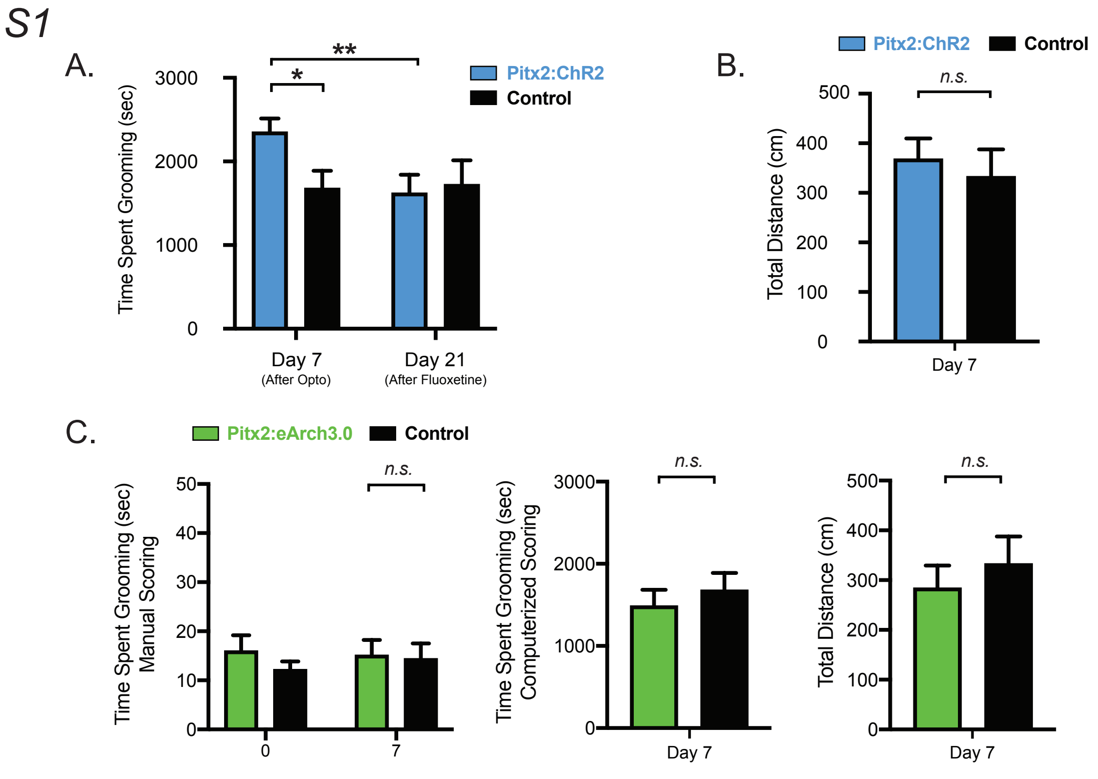

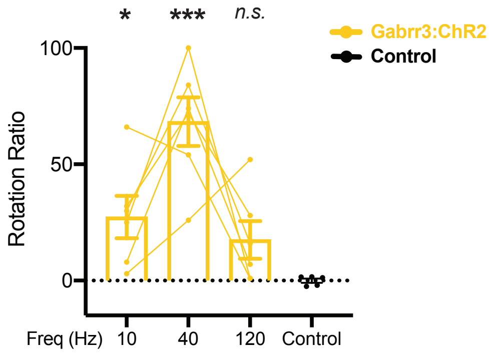

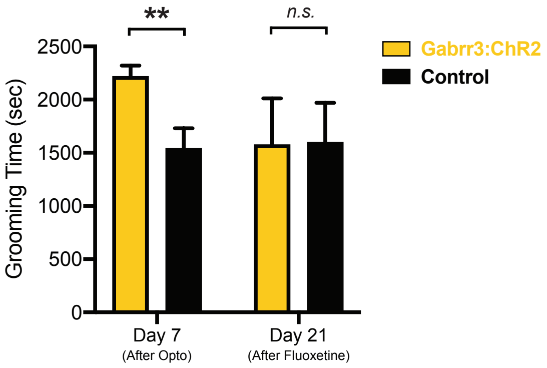

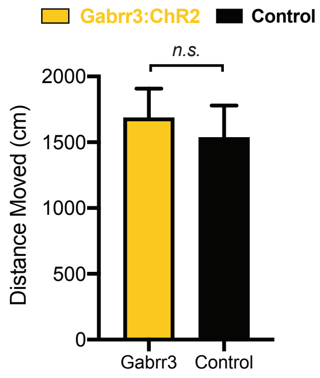
